# Host retargeting predominated over gene transfer during early chromatophore integration in *Paulinella micropora*

**DOI:** 10.64898/2026.04.14.718432

**Authors:** Moisès Bernabeu, Toni Gabaldón

## Abstract

The relative contributions of gene acquisition and protein retargeting to early organellogenesis remain debated. We addressed this question for the recently evolved chromatophore of *Paulinella micropora*, which provides a unique window into the initial stages of primary organelle integration. Using a stringent phylogenomic pipeline, we identified 282 *P. micropora* gene families whose origin can be attributed to horizontal gene transfer (HGT) with high confidence, including 115 for which we could confidently assess the donor lineages. In addition to the expected cyanobacterial signal from the chromatophore ancestor, we detected recurrent contributions from Gammaproteobacteria, Bacteroidota, Actinobacteriota, and Chlamydiota. Relative stem-length analyses indicate that these donors contributed in temporally structured periods (waves), with several non-cyanobacterial signals older than the median cyanobacterial signal and others clearly more recent. HGT-derived genes are enriched in transport and metabolic functions, which might have been relevant to host-symbiont interaction. However, based on computational predictions, the chromatophore-targeted proteome is dominated by vertically inherited host genes with only 36 of 287 predicted chromatophore-targeted nucleus-encoded proteins being of HGT origin, the majority of which are absent in *Paulinella chromatophora*, suggesting a recent origin in *P. micropora*lineage. Thus, although repeated horizontal acquisition from multiple bacterial partners shaped the nuclear genome of *P. micropora*, early chromatophore integration appears to have been dominated by host-derived retargeting, with HGT providing a smaller but detectable complementary contribution. These results support a mixed model of organellogenesis in which targeting and gene transfer act together, but with host retargeting playing the dominant early role.

## Introduction

The origin of new organelles through endosymbiosis is a rare but transformative event that has had profound impacts in the evolution of eukaryotes (Archibald 2015). Two such events underlie major innovations in eukaryotes: the acquisition of an alpha-proteobacterial endosymbiont that gave rise to mitochondria prior to the diversification of all known eukaryotic lineages (López-García and Moreira 2023), and the engulfment of a cyanobacterium that originated the plastid and conferred photosynthetic capacity to eukaryotes (Parfrey et al. 2011; de Vries et al. 2016). Photosynthesis later spread to other groups through secondary endosymbioses (Keeling 2013; Sibbald and Archibald 2020).

Central to the organellogenesis process is the emergence of host-endosymbiont interdependence, a process that is strengthened by horizontal gene transfer (HGT) from the symbiont to the host nucleus (sometimes termed endosymbiotic gene transfer, EGT), and by the evolution of targeting systems that import nuclear-encoded proteins back into the nascent organelle (Gabaldón and Pittis 2015). However, the relative order and contributions of these events during the earliest steps of organellogenesis remain debated (Keeling 2024). Competing hypotheses propose that organelle integration is initiated either by gene acquisition from the symbiont (an “HGT/EGT-first” view) or by the early establishment of protein targeting (a “targeting-first” view).

The photosynthetic organelle in autotrophic *Paulinella* species, the chromatophore, represents a uniquely recent case of primary photosynthetic organelle origin besides the canonical chloroplast (Marin et al. 2005), offering an exceptional window into early events of organellogenesis (Gabr et al. 2020). The chromatophore derives from a single endosymbiotic event involving a Cyanobacterium of the Prochlorococcus-Synecoccocus clade (Marin et al. 2005), estimated to have occurred around 90-140 Mya (Delaye et al. 2016). Because this endosymbiotic event is far more recent than the origin of chloroplasts or mitochondria, *Paulinella* offers a unique model to examine evolutionary patterns that are largely obscured in older organelles.

Previous analyses of autotrophic *Paulinella* nuclear genomes have revealed a mosaic of genes with non-vertical origins, including contributions from the chromatophore ancestor and other cyanobacterial, alphaproteobacterial and gammaproteobacterial clades (Nowack et al. 2016; Lhee et al. 2021). In this context, Nowack *et al* . (2016) suggested the possibility of HGT of post-symbiotic bacteria to the eukaryote. However, key questions remain unresolved. For instance, what is the relative temporal order of these transfers, do they occur in discrete waves from particular donors, and how did transferred genes contribute to establishing host-chromatophore integration? Finally, previous studies have not addressed the question of the relative tempo and relevance of gene transfer and protein retargeting in early chromatophore integration.

In this study, we generated a timeline of HGT in the *Paulinella micropora* nuclear genome, the most complete and contiguous genome assembly and proteins set for *Paulinella*, using a stringent, phylogeny-informed, pipeline to identify high-confidence HGT events, infer their likely bacterial donors, and estimate their relative ages. We then relate phylogenetic origin to predicted function and subcellular localisation, with a particular focus on the imported chromatophore proteome. Our results point to multiple significant bacterial donor lineages beyond the cyanobacterial ancestor of the chromatophore and identify transfer waves of differing intensity that likely span pre-, peri-, and post-endosymbiotic phases, including signals consistent with a close pre-chromatophore association between *Paulinella* and Burkholderiales-like bacteria. Importantly, we find that chromatophore-targeted proteins are dominated by vertically inherited host genes, with a smaller but detectable contribution from HGT-derived genes of diverse bacterial origins. Taken together, these results support a “mixed ratchet” view of organellogenesis in which early targeting-mediated integration and EGT processes jointly drive the chromatophore along an irreversible path toward organelle status.

## Results and discussion

### Diverse origins and functions of HGT genes in P. micropora

To identify HGT events from prokaryotes into the nuclear genome of *P. micropora*, we first clustered *P. micropora* proteins into families and searched for homologues in a custom database spanning diverse prokaryotic, eukaryotic and viral sequences (see Materials and Methods). We then applied a step-wise pipeline combining an initial blast-based approach and followed by a phylogeny-based analysis to define three nested sets of HGT candidates with increasing stringency. As an initial screen, we used a blast-only approach to select families whose best hit was non-eukaryotic (i.e. from prokaryotes or viruses), or for which more than 50% of the top-1000 blast hits were non-eukaryotic. This permissive filter identified 2,709 genes in 2,623 putative HGT events (post-HGT gene duplication results in multiple genes per event), hereafter referred to as HGT precandidates (Figure 1a). This set is likely to comprise bona-fide HGT cases as well as many vertically inherited genes with widely distributed homologues, and we only used it as a basis for further exploration (Supplementary Table 1).

**Figure 1.**
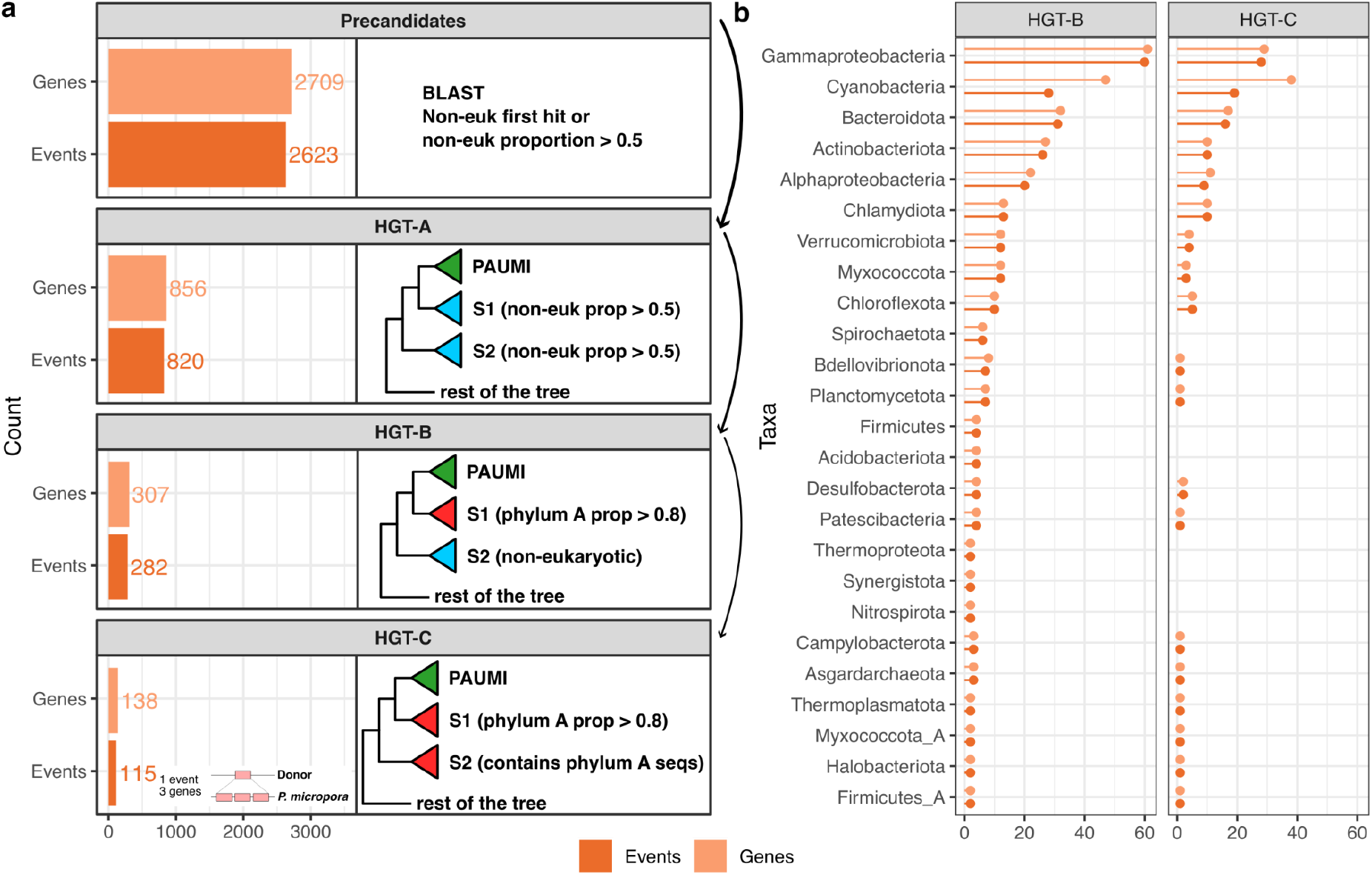
Horizontal gene transfers in *Paulinella micropora* . a) Step-wise selection of HGT candidates, from less-stringent (top) to more stringent (bottom) filtering steps. Each box corresponds to one nested step and within each box, left bars indicate the number of genes and events (i.e. families) selected, whereas the text or scheme on the right indicates BLAST or tree topology filtering criteria. Lighter colour bars show the number of events (number of transfers from the donor to the acceptor), while darker colour bars show the number of genes (the total count of genes transfers, accounting for posterior expansions after the transfer event). The box in HGT-C shows the difference between number of events and number of genes due to post-HGT expansions. b) Number of genes and events assigned to the different prokaryotic taxa in the HGT-B and HGT-C sets.

We next used a fast phylogenetic reconstruction approach on each HGT precandidate and its 1,000 closest homologs, and assessed the resulting topology as an additional filter (see Materials and Methods). HGT precandidates that had a first and second sister clades containing more than 50% non-eukaryotic sequences were further considered as potential HGT candidates (Figure 1a). This set (HGT-A), comprising 820 families (856 genes), is likely to be enriched in bona-fide HGT genes, but may still contain some vertically inherited genes or genes acquired before the origin of the *Paulinella* genus with dubious phylogenies. To obtain a more reliable subset of HGT candidates (HGT-B) we selected up to 300 homologs that were topologically closest to each HGT-A family in the respective phylogeny, and reconstructed a new phylogeny using a more accurate approach (see Materials and Methods). We selected a family as HGT-B when the first sister clade contained more than 80% sequences from the same non-eukaryotic lineage, the second sister clade contained a majority of non-eukaryotic sequences, and the branch subtending the *P. micropora* family and its sister group had bootstrap support greater than 80%. This procedure defined 282 HGT-B families (307 genes) with a well-supported HGT topology and a clearly identified sister-clade (Figure 1a). In a subset of these candidates (HGT-C, n = 115 families), the most abundant phylum within the first sister clade was also found in the second sister clade, providing stronger support to the putative taxonomic identity of the prokaryotic donor (Figure 1a, strict). These HGT-C families comprised 138 genes with a well-supported HGT origin and a well-characterized putative donor. Previous studies using more relaxed thresholds identified 148 and 229 genes with bacterial ancestry in *P. micropora* (Lhee et al. 2021) and *P. chromatophora* (Nowack et al. 2016), respectively. These differences may result from a higher coverage of prokaryotic diversity in our study.

The phylogenetic analysis identified 25 putative gene donors at the phylum level in HGT-B, 19 of which were also found in the HGT-C subset. Consistent with the known cyanobacterial origin of the chromatophore, the relative contribution of cyanobacterial genes increased in the more stringent subset from 9.93% (HGT-B) to 16.5% (HGT-C). Additionally, as a positive control of our donor identification procedure, we run our HGT detection pipeline on the chromatophore genome (see Methods and Supplementary Discussion). Of the 859 chromatophore-encoded proteins, 796 passed the HGT-C filters, with most of them (791) being assigned to Cyanobacteria. Altogether, these results support the ability of our pipeline to recover the expected cyanobacterial ancestry in a control dataset of known origin. Only five chromatophore genes were assigned to non-cyanobacterial donors in the HGT-C set. This indicates that, although the cyanobacterial ancestor of the chromatophore may have exchanged genes with other non-cyanobacterial prokaryotes prior to the endosymbiosis, this non-cyanobacterial signal remains minimal and cannot drive the non-cyanobacterial origins of nuclear HGTs. Based on this analysis, we concluded that the HGT-C set is a good proxy for HGT-derived *P. micropora* genes and their donors.

Interestingly, the results for the nuclear genome show a diversity of significant donors beyond the cyanobacterial ancestor of the chromatophore (Figure 1b). Because some assignments may reflect phylogenetic uncertainty, we focused on phyla involved in at least 10 transfer events, as consistent signals are less likely to stem from stochastic errors. Gammaproteobacteria appears as the most frequent donor in the HGT-C subset in terms of number of events (28), followed by Cyanobacteria (19), Bacteroidota (16), Actinobacteria (10), Chlamydia (10), 14 additional potential donors were involved in up to nine events each. Some HGT events were followed by duplications, thereby increasing the number of HGT-derived proteins beyond the number of HGT events. The most notable example is the high light-inducible (*hli*) protein family, which expanded to 20 in-paralogs and has been previously studied in *P. chromatophora* (Calatrava et al. 2022).

Together, our analyses reveal that the *P. micropora* nuclear genome bears a diverse signature of HGT from bacterial donors beyond the chromatophore ancestor. As compared to a previous analysis of the *P. chromatophora* genome, which identified eight bacterial donor clades: Proteobacteria, Cyanobacteria, Actinobacteria, Aquificae, Chlamidiae/Verrucomicrobia, Chloroflexi, Firmicutes and Panctomycetota (Nowack et al. 2016), our results expand the list of donors in the case of *P. micropora* using more stringent criteria and a taxonomically broader dataset of prokaryotes.

### Distinct gene transfer waves reveal five major bacterial contributors

We next focused on the five donors that contributed at least 10 events in the high confidence HGT-C set. Recurrent transfers from the same donor may reflect sustained ecological associations between the donor and the recipient species, raising the possibility that *Paullinela* maintained stable interactions with bacterial partners beyond the endosymbiont that originated the chromatophore. To place these interactions in a relative temporal framework, we estimated the timing of gene acquisitions using a branch length ratio approach (see Methods, (Pittis and Gabaldón 2016; Bernabeu et al. 2025)). Specifically, for each HGT event, we measured the length of the branch connecting the *P. micropora* gene (or the node representing the last common ancestor of the gene family) to its first sister clade. This ‘stem length’ represents the sequence divergence from the ancestral transferred sequence (the first node of the *P. micropora* family). To account for evolutionary rate variation across gene families, we normalized this length by the median root-to-tip distance for the corresponding phylogeny. This yielded distributions of relative transfer ages for each donor.

Based on the medians of these distributions, we inferred the following relative order among the major donors, from the oldest (longer stem lengths) to the most recent (shorter stem lengths): Bacteroidota, Actinobacteriota, Gammaproteobacteria, Cyanobacteria and Chlamydiota (Figure 2a). Notably, Actinobacteriota and Cyanobacteria show narrower distributions than the remaining donors (particularly Gammaproteobacteria), which may reflect more episodic events rather than a continuum of transfers (Figure 2b). Because there is no complete genome from heterotrophic *Paulinella* species and the representation of autotrophic *Paulinella* species is still limited, we cannot accurately determine whether HGT from non-cyanobacterial donors predated or followed chromatophore acquisition. However, most non-cyanobacterial donors exhibit longer median stem lengths, suggesting earlier interactions. Conversely, we identified transfers that are younger than those from Cyanobacteria, indicating that the *P. micropora* genome is still dynamic with regard to HGT and has acquired genes from other donors after the establishment of the chromatophore. Additionally, we found 36.23% HGT-C families (50 out of 138) in *P. micropora* with no homologues in *P. chromatophora* . Their absence from *P. chrompatophora*, supports that HGT continued after the speciation from the two autotrophic species, and therefore, after the chromatophore acquisition (Nowack et al. 2016; Lhee et al. 2021).

**Figure 2.**
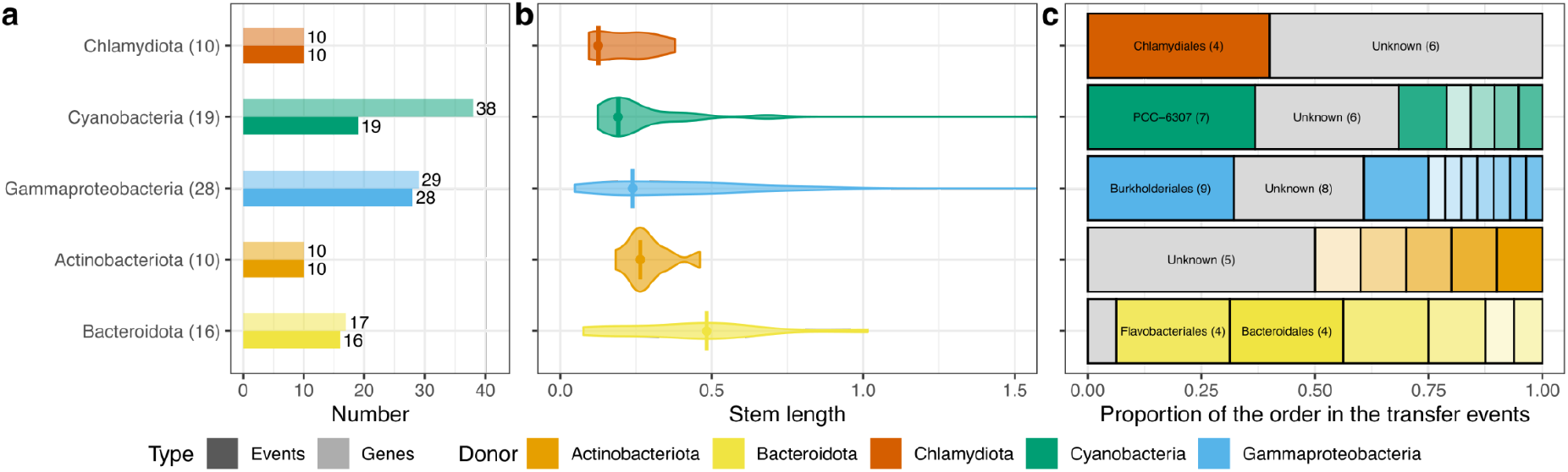
Relative timing of the gene transfers to *P. micropora* . a) Number of events (darker color) and genes (lighter color) from each donor, indicated on the Y axis and colored as in the legend. b) Distributions of normalised stem lengths for protein families assigned to each donor. c) Proportion of events assigned to each donor at the order level. Grey color indicates unknown donor at the order level.

To infer the function of the *P. micropora* proteins, we used an in-house annotation pipeline that integrates complementary strategies (see Supplementary Methods). This pipeline annotated 15,404 out of 32,361 proteins (Figure 3a, Supplementary Table 1), substantially improving on the 11,493 proteins previously annotated by Lhee et al. (2021). Notably, our annotated dataset included 298 of the 307 HGT-B candidates (Figure 3b). We next examined the functional annotations of the transferred proteins. Genes involved in carbohydrate and amino acid metabolism were often acquired from Bacteroidota and Actinobacteria; genes involved in nucleotide transport, post-translational modifications and cell wall and envelope biogenesis from Bacteroidota; genes related to energy production and conversion from Cyanobacteria; and genes involved in DNA processing and cell cycle were predominantly acquired from Gammaproteobacteria (Supplementary Figure 1).

**Figure 3.**
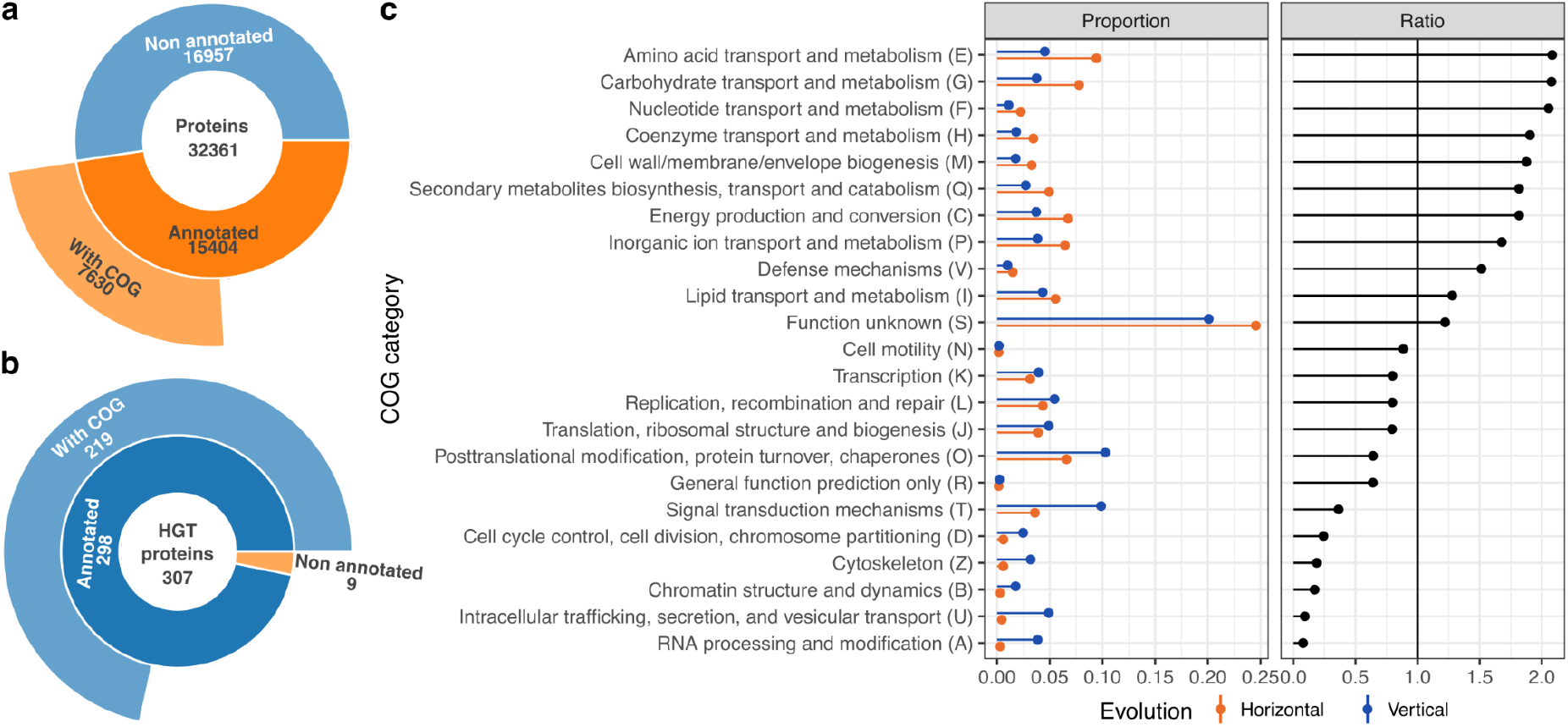
Annotation summary of the *P. micropora* proteins. a) Number of proteins and COG categories annotated in the whole genome. b) Number of proteins and COG categories annotated in the subset of HGT genes. c) The proportion of each COG category in the HGT and vertical genes (coloured as in the legend), together with their proportion ratio. Values above 1 (black solid line) indicate categories overrepresented among HGT genes relative to vertically inherited genes.

To define the taxonomic affiliation of the donors at a higher resolution, we calculated the proportion of events at the order level (Figure 2c). Importantly, we detect seven *Prochlorococcus* – *Synecocchocus* (PCC-6307) transfers out of the 19 cyanobacterial transfers (37%). This proportion appears unexpectedly low, as it is the sister clade to the chromatophore (Supplementary Figures 2 and 3). Limited phylogenetic resolution at the order level likely contributes to this pattern. Supporting this interpretation, only 591 out of 859 (69%) of the chromatophore-encoded proteins are inferred to have this ancestry. Comparable uncertainty at the order level should therefore be assumed for other donors. Additionally, transferred genes may retain weaker phylogenetic signals as compared to genes still retained in the chromatophore, because of post-HGT sequence amelioration. Strikingly, the number of transfer events assigned to Burkholderiales was comparable to that assigned to Cyanobacteria. This suggests a close interaction between a species within Burkholderiales and *Paulinella*, perhaps through the establishment of an ancestral symbiotic interaction, or, alternatively, through common environmental co-occurrence. HGT from Burkholderiales have been previously identified in *P. chromatophora* (Nowack et al. 2016), supporting an early interaction with the *Paulinella* autotrophic lineage as inferred by our relative dating. Further assessing the nature and timing of the potential *Paulinella-* Burkholderiales association would require broader genome sampling within and around the *Paulinella* genus, as well as additional characterization integrating cell biology and ecology.

Functional profiling shows that transport and metabolism-related categories are enriched among HGT-B genes (which are reliable HGTs, independently of the certainty on the donor), whereas informational genes dominate among vertically-evolving genes. In particular, functions related to amino acid, carbohydrate, nucleotide, and coenzyme transport and metabolism are approximately twofold more frequent among HGT-B genes than among vertically-evolving genes (Figure 3c). Interestingly, the category “energy production and conversion” is enriched in HGT-B, by approximately 1.5, and includes genes from at least six bacterial phyla, including Gammaproteobacteria and Cyanobacteria. Together, these results indicate that HGT has made substantial contributions to the metabolic repertoire of *P. micropora* . Moreover, the enrichment in transport-related functions suggests a potential contribution of transferred genes from various donors to the establishment and diversification of host-chromatophore interactions.

### Vertically-inherited proteins dominate the chromatophore-targeted proteome

We inferred the subcellular targeting of nuclear-encoded genes using DeepLoc, PrediSi and SignalP. Because of its training design, DeepLoc does not recognize the chromatophore as a possible localisation. Moreover, it assigned 139 proteins to the plastid, an organelle absent from *P. micropora* . These plastid predictions showed little overlap with the 287 chromatophore-targeted proteins identified by Lhee et al. (2021) using HMM profiles of the chromatophore transit peptide (crTP) (21 shared proteins). We therefore discarded DeepLoc plastid predictions and used only the crTP-based set as our best proxy for the chromatophore-targeted proteome, despite they do not cover the full set of experimentally-detected targeted proteins (Singer et al. (2017), and Supplementary Discussion). We kept DeepLoc predictions for the remaining localizations. Relative to vertically-inherited, HGT proteins were more frequently predicted to be secreted extracellularly or targeted to the peroxisome, chromatophore and mitochondria. In contrast, vertically-inherited proteins were more frequently predicted to locate in the cytoplasm, cell membrane, and nucleus (Figure 4a).

**Figure 4.**
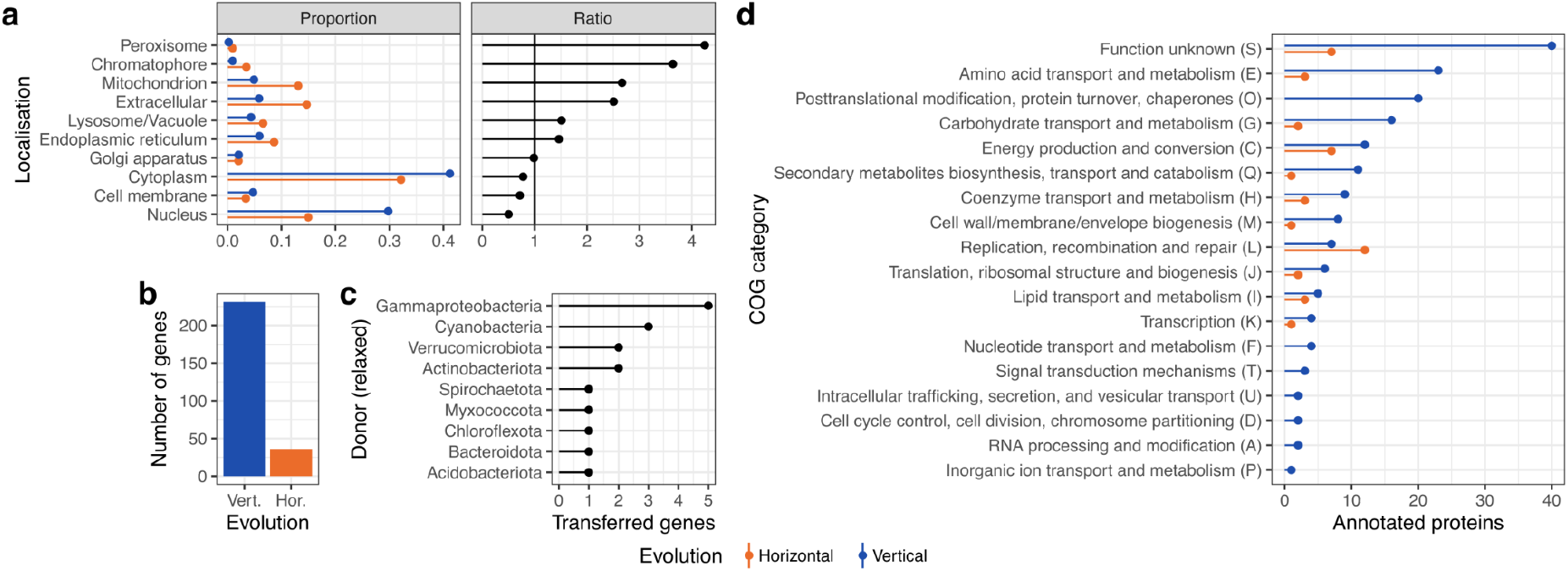
Subcellular localisation of the *P. micropora* proteins. a) Proportion of proteins from each of the inferred origins that are assigned to each subcellular compartment, and the corresponding proportion ratio. Ratio values greater than 1 indicate that the predicted localisation is more frequent among HGT-derived proteins than among vertically-inherited proteins. b) Number of nuclear-encoded proteins predicted to be targeted to the chromatophore based on the presence of a chromatophore transit peptide (crTP) as inferred by Lhee et al. (2021), grouped by inferred origin (vertical or horizontal). c) Putative donor of the HGT-derived proteins predicted to be targeted to the chromatophore. d) Functional categories of chromatophore-targeted proteins according to inferred origin.

Among the nucleus-encoded proteins predicted to be targeted to the chromatophore, only 36 (12.54%) were classified as HGT-B (Figures 4b and 5a), indicating that most of the chromatophore’s imported proteome is of vertical origin, consistent with previous reports (Nowack et al. 2016; Lhee et al. 2021). However, when compared with the overall representation of HGT-B in the nuclear genome (307/32361, 0.9%) it can be concluded that proteins of HGT origin are over-represented in the chromatophore. Despite their low overall number, these HGT proteins were inferred to derive from nine putative donors at the phylum level. Importantly, Gammaproteobacteria was the most common donor with five HGTs, surpassing the chromatophore-ancestor Cyanobacteria, which contributed only three (Figure 4c). Despite these results, Cyanobacteria is the most prevalent taxon when including a broader set of proteins imported into the chromatophore using homologues to the small chromatophore targeted proteins (sCTPs) in *P. chromatophora* (see Supplementary Discussion, Supplementary Table 2, and Supplementary Figure 4). These results suggest that piecemeal retargeting of HGT genes from various origins, and not only from the chromatophore ancestor, has shaped this organelle’s proteome.

We compared the 36 chromatophore-targeted HGT proteins to the *P. chromatophora* predicted nuclear-encoded proteome and found 20 putative homologs, of which only two had cyanobacterial origins. In contrast, out of the 251 chromatophore-targeted proteins of vertical-inheritance, we could find *P. chromatophora* homologs for 112. Although some missing hits might be explained by the highly incomplete or fragmented annotation of the *P. chromatophora’s* proteome (Nowack et al. 2016), our observation suggests that retargeting of proteins from various origins occurred both before and after the *P. micropora* / *P. chromatophora* speciation, and may still be ongoing. Also, these numbers suggest a similar fraction of pre-speciation retargeting for HGT proteins of cyanobacterial (2/3, 66%) or non-cyanobacterial origins (20/33, 61%), and a slightly lower one for vertically-inherited proteins (112/251, 45%). Importantly, although proteins of cyanobacterial origin are more likely to predate this speciation than vertically inherited proteins, they represent a very small portion of the pre-speciation imported proteome (2/133, 1.5%). This scarcity suggests that protein retargeting from non-EGT sources dominated the early steps of chromatophore evolution (Figure 5b).

**Figure 5.**
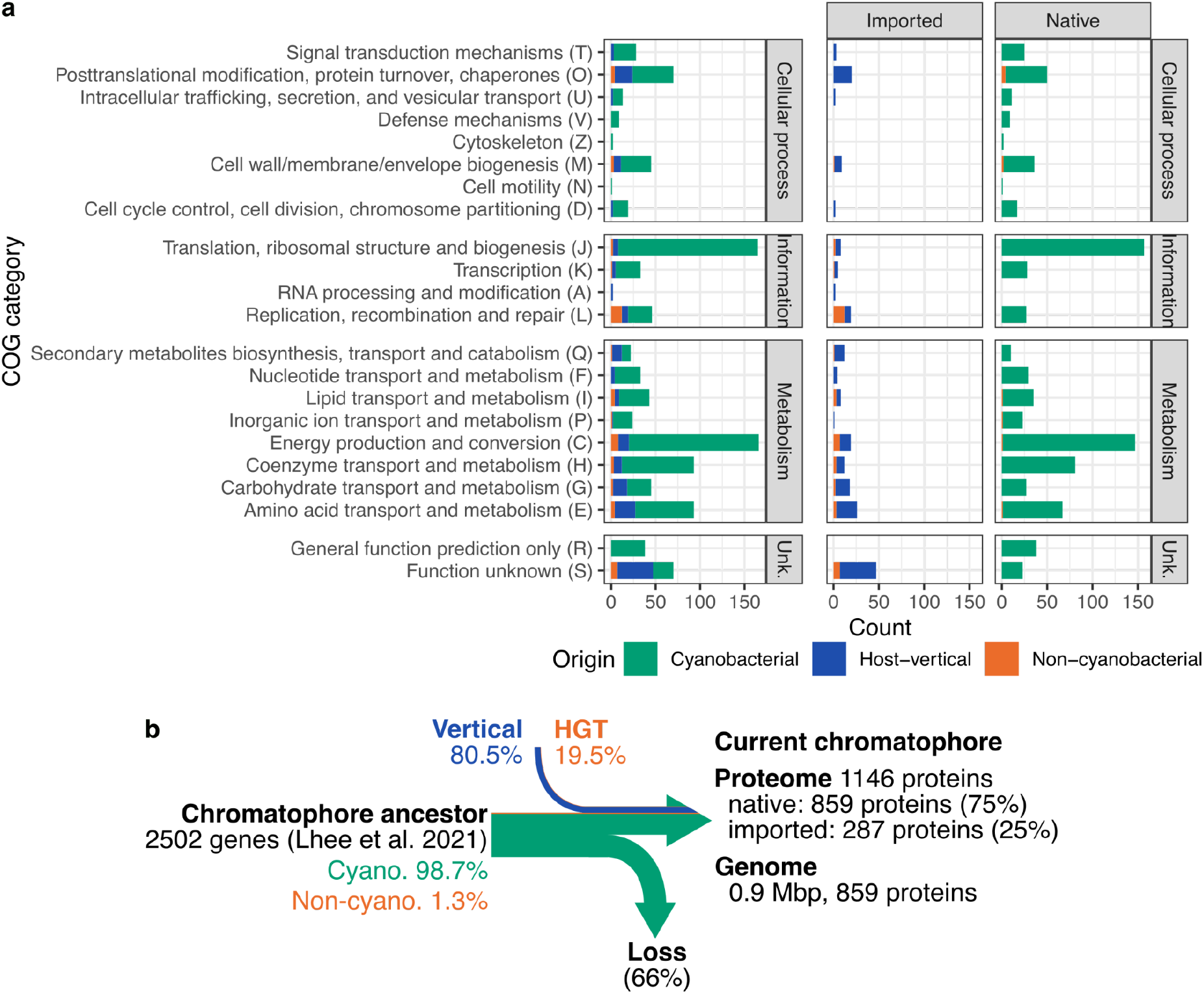
Origins of the chromatophore proteome. a) Count of proteins from a certain origin annotated with the functional category of the y-axis for the full proteome (left), for the chromatophore-encoded proteins (center), for the host encoded-proteins (right). b) Overview of the contribution to the chromatophore proteome of *P. micropora* .

The most represented functions among vertically-inherited proteins in chromatophore-targeted proteome were related to transport and metabolism (48, 16%), post-translational modification (20, 7%), and energy production (12, 4%) (Figures 4d and 5a). This means that the vertically inherited proteins targeted to the chromatophore contribute to central functions of the chromatophore. We inspected specific metabolic functions of vertically-inherited proteins retargeted to the chromatophore (Supplementary Table 3), and found that many of them replace functions of genes that are inferred to be present in the ancestor of the chromatophore (see Supplementary Discussion, and Supplementary Figure 5). Such replacement could have triggered the dependence of the symbiont on the host for key metabolic processes such as the synthesis of amino acids and nucleotides.

HGT-derived imported proteins contribute mostly to replication, recombination and repair (12/36, 33%), as well as energy production (7/36, 19%). Notably, HGTs from multiple bacterial donors contributed to functions related to informational metabolism, as previously reported for *P. chromatophora* (Nowack et al. 2016; Singer et al. 2017). However, unlike vertically inherited enzyme-coding genes, chromatophore-targeted enzymes of HGT origin perform rather peripheral and isolated reactions in the metabolism of the chromatophore (see Supplementary Discussion and Supplementary Figure 5). Taken together, these patterns suggest that components of the chromatophore informational machinery were acquired from different bacterial donors, integrated into the host nuclear genome, and subsequently, or concurrently, retargeted to the chromatophore. Such retargeting may have been favoured because bacterial-derived and not eukaryotic components were more readily integrated into a bacterial-like system. They could either reinforce the host-symbiont interdependence, but, as their role in metabolic pathways they could have broadened the metabolic capacities of the nascent organelle. As for the three imported proteins from cyanobacterial origin, they all have annotated functions likely related to secondary metabolism, including GTPase, a 3-(3-hydroxy-phenyl)propionate hydroxylase, and a CMP-N-acetylneuraminate monooxygenase, with the latter one being retargeted post-speciation.

Overall, the localisation, origin, and functional composition of chromatophore-targeted proteins support a clear predominance of host-derived genes (Singer et al. 2017), in agreement with previous work (Lhee et al. 2021). When considered alongside current models of EGT (Keeling 2024), our results suggest that the retargeting of nuclear-encoded proteins (either from vertical or horizontal origins) occurred much more frequently than EGT in the early phases of chromatophore integration. Proteins, regardless of their evolutionary origin, may have been retargeted to the cyanobacterial endosymbiont, and those that improved communication and metabolic integration between host and symbiont were subsequently retained, making the association increasingly irreversible.

This scenario is consistent with the “targeting ratchet” model (Keeling 2013), which has also been proposed for the kleptoplast of *Rapaza viridis* (Karnkowska et al. 2023). Overdominance of host-derived proteins, particularly in categories related to core regulatory functions and metabolite transport, had also been observed for the mitochondrion, where only 16% of the proteome can be traced back to an alphaproteobacterial origin (Gabaldón and Huynen 2003; Gabaldón and Huynen 2007). Our results, for a more recent organelle, suggest that this predominance of host-derived proteins starts very early during organellogenesis. They also indicate that import of host-derived proteins, more than HGT as proposed in the HGT ratched model (Nowack et al. 2016), is the primary process shaping the organellar proteome and compensating the progressive reduction of the organellar genome. Finally, the functions of the retargeted proteins of cyanobacterial origin seem to be involved in chromatophore-specific functions rather than in host-symbiont integration.

## Conclusions

The chromatophore of autotrophic *Paulinella* species provides a uniquely recent model to investigate the early stages of symbiotic organellogenesis (Nowack and Weber 2018; Coale et al. 2024). By combining stringent phylogenetic analysis, relative dating, with functional and targeting prediction, we show that the proteome of *P. micropora* has been shaped by continuous HGTs from various donors beyond the cyanobacterial ancestor of the chromatophore. These transfers were recurrent and temporally structured, indicating that chromatophore evolution took place in the context of broader and sustained interactions with diverse bacterial partners. For instance, our analysis suggests a close interaction between *P. micropora* and a bacterium from Burkholderiales, resulting in a wave of transfers shortly predating the transfers from the cyanobacterial endosymbiont.

Moreover, our results indicate that retargeting of nuclear-encoded proteins to the chromatophore is an ongoing process that involves genes of different origins. Importantly, our results depict an early chromatophore-imported proteome dominated by host-derived proteins that frequently substitute enzymes likely present in the ancestor of the chromatophore. Additionally, the targeted genes include only a few proteins of cyanobacterial origin and a surprisingly higher number of HGT-derived proteins from other donors, that play a rather peripheral role in the metabolic network of the chromatophore. However, these inferences may be incomplete due to the lack of subcellular localisation studies in *P. micropora* .

Taken together, these results support a “mixed ratchet” model in which organellogenesis proceeds through the combined action of the targeting ratchet (Keeling 2013) and endosymbiotic ratchet (Bhattacharya et al. 2023), with an earlier role of retargeting of host-derived proteins. In this sense, the data are consistent with a scenario in which targeting ratchet-like processes began early during host–symbiont integration, whereas endosymbiotic ratchet-like effects may have become more important as the association became increasingly irreversible. Although this framework provides a plausible explanation for the mosaic origin and timing of chromatophore-associated genes, the precise population-genetic and ecological sequence of these events remains to be tested. More broadly, our findings suggest that early organellogenesis does not require extensive initial gene transfer, but can instead be driven primarily by host retargeting, with HGT from various donors providing a secondary and complementary source of innovation.

## Materials and Methods

### Paulinella micropora genome and database configuration

We downloaded the genome assembly of *Paulinella micropora* KR01 (Lhee et al. 2021) from the Mycocosm database (JGI, https://mycocosm.jgi.doe.gov/Paumic1/Paumic1.home.html, accessed 4 December 2023). We assessed genome completeness and contamination using OMArk v0.3.1 (Nevers et al. 2025), and detected no contaminant proteins (Supplementary Tables 4 and 5).

We initially annotated the function of *P. micropora* proteins using EggNOGmapper v2.1.12 (Cantalapiedra et al. 2021) and InterProScan v5.57-90.0 (Jones et al. 2014). As nearly 60% of the proteins remained unannotated, we additionally performed profile-based searches with hmmsearch v3.3.2 (Eddy 2011) against the Database of Clusters of Orthologous Genes (COG) v2020 (Galperin et al. 2015), PFAM v35.0 (Mistry et al. 2021), KEGG Orthology (KO) v2023-02-01 (Kanehisa et al. 2016) and TIGR (Haft et al. 2013), accessed on 20 April 2023. We filtered the HMM results to keep the first hit and merged these annotations together with those coming from the EggNOGmapper and InterProScan predictions explained above, using a custom python script that produced a single annotation file (see Supplementary Methods, Supplementary Figure 6 and Data and Code availability for details). Additionally, we added localisation and signalling data, as inferred with DeepLoc v2 (Thumuluri et al. 2022), SignalP v5.0b (Almagro Armenteros et al. 2019) and PrediSi (Hiller et al. 2004), this last one ran on the web server (http://www.predisi.de/index.html, accessed in February 2024).

For subsequent comparative analyses, we assembled a custom database comprising protein sequences from different curated databases, aiming to cover most of the known diversity. In brief, we collected archaeal and bacterial proteins from GTDB v207 (Parks et al. 2020), all eukaryotic reference genomes from UniProt v2023_v1 (The UniProt Consortium 2023), as well as the clustered version of the Reference Viral DataBase RvDB v26.0 (Goodacre et al. 2018). We further populated this database for the Rhizaria clade using 15 available genomes previously used in van Hooff and Eme (2023). We taxonomically annotated this custom database with the taxonomic lineage file (see Supplementary Material), which was generated using UniEuk taxonomy for eukaryotes, GTDB taxonomy for archaea and bacteria and RvDB taxonomy for viruses. We next inferred *P. micropora* protein families by clustering them with mmseqs2 release 14-7e284 (Steinegger and Söding 2017) easy-linclust algorithm using stringent thresholds: 90% minimum identity and 80% minimum coverage (-c 0.8 --min-seq-id 0.9). This resulted in 590 protein clusters that comprised 1,349 proteins, of which the largest cluster contained 20 proteins, and 31,012 singletons.

### Inferring HGT candidates

To infer HGT candidates, we first searched homologs for each of the annotated *P. micropora* proteins in the custom database explained above, by using BLAST v2.13 (Altschul et al. 1990). We established a minimum e-value threshold of 1e-5 and a minimum alignment coverage over the query sequence of 75%. We further removed all hits that were either 2.5 times longer or 2.5 times shorter than the query to avoid hits driven only by local homology restricted to a single domain. After filtering, we concatenated the unique BLAST results for all queries belonging to the same cluster (family), and selected up to 1,000 hits with the lowest e-value. Those *P. micropora* families and proteins for which the proportion of non-eukaryotic hits in the top-1000 BLAST results was higher than 50% or those that had a non-eukaryotic sequence as first hit were considered “ *HGT precandidates”* .

For each HGT pre-candidate protein or family, and their selected BLAST hits, we reconstructed a phylogeny using a fast approach. For this, we aligned the sequences using FAMSA v2.1.2 (Deorowicz et al. 2016), trimmed with trimal v1.4.rev15 (Capella-Gutierrez, Silla-Martineofand Gabaldón, 2009) using the -gappyout option, and reconstructed a tree with FastTreeMP v2.1.11 (Price et al. 2010) under the LG+G4 model (Yang 1994; Le and Gascuel 2008). We rooted and scanned the trees using ETE3 (Huerta-Cepas et al. 2016). As a root, we used the non-eukaryotic sequence in the tree with maximum phylogenetic distance to the *P. micropora* protein or family. This rooting provides a tree in which the *P. micropora* sequences are far from the root, providing directionality to its phylogenetic neighbourhood. We then assessed the taxonomy of the sequences contained in the two closest sister clades to the *P. micropora* proteins (which were always monophyletic in our trees). Pre-candidate proteins or clusters for which each of the two sister clades contained more than 50% non-eukaryotic sequences were considered *HGT candidates* (HGT-A).

To infer a more stringent subset of HGT proteins, we reconstructed a phylogeny of the HGT-A candidates using a more accurate phylogenetic reconstruction: the phylomeDB pipeline (Fuentes et al. 2022) with some modifications. In brief, we filtered the 300 first hits of each HGT candidate and aligned them in forward and reverse orientation using three different aligners: MAFFT v7.490 (Katoh and Standley 2013), muscle5 v5.1 (Edgar 2022) and kalign v3.3.2 (Lassmann 2020). The resulting six alignments were merged using MergeAlign (Collingridge and Kelly 2012), which assesses the column precision and calculates a consensus alignment. This alignment was then trimmed using trimAl v1.4.rev15 (Capella-Gutierrez, Silla-Martineofand Gabaldón, 2009) using the -gappyout option. We reconstructed the phylogeny using IQ-TREE v2.2.0 (Minh et al. 2020) under the fittest model according to ModelFinder (Kalyaanamoorthy et al. 2017). We used a subset of the available models to speed up the selection, including single-matrix models (WAG (Whelan and Goldman 2001), LG (Le and Gascuel 2008), JTTDCMut (Kosiol and Goldman 2005) and JTT (Jones et al. 1992) and multiple matrices models (LG4X and LG4M), which account for a certain degree of exchangeabilities matrix heterogeneity (Le et al. 2012). We assessed branch support using 1,000 ultra-fast bootstrap replicates (Hoang et al. 2018). Finally, we rooted the trees at the farthest non-eukaryotic sequence to the *P. micropora* family or sequence.

Using the trees of the HGT-A set, we analysed the taxonomic affiliation of the first and second sister clades to the *P. micropora* protein family. To be considered HGT-B, we required the HGT-A family to be nested within at least two subsequent clades, each with a minimum of 66% non-eukaryotic sequences. We also established a minimum support consisting of an ultrafast bootstrap threshold of 80% in the branch connecting the *P. micropora* family with its first sister (i.e. support value for the acquisition). This nestedness within non-eukaryotic sequences prevents us to misdetect eukaryote-to-prokaryote HGTs as prokaryote-to-eukaryote ones. Finally, among these HGT-B genes, we defined as HGT-C, as those HGT-B families, where the *P. micropora* proteins were nested within a specific bacterial phylum, requiring both sisters to have at least 80% sequences from the same prokaryotic phylum each. To reinforce the certainty of these observations, we used an ultrafast bootstrap threshold of 85% for the branch subtending the two sister clades. And assessed the robustness of the donor using a “taxonomic bootstrap” approach (see Supplementary Results and Supplementary Figure 7).

The only available peptide set for of *P. chromatophora* is a highly fragmented transcriptome-derived proteome (Nowack et al. 2016). We attempted to use this dataset, but its fragmentation resulted in difficulty of establishing homology relationships (Supplementary Results, and Supplementary Figure 8a) and generated gap-rich alignments (even in BUSCO markers) that would result in unreliable phylogenetic trees (Supplementary Figure 8b). Hence, we decided not to include these proteins in the phylogenetic analyses but only use them to assess the potential presence of *P. micropora* homologues in *P. chromatophora* . For this, we performed a BLAST search, and filtered the results using a minimum identity and coverage of 50%, and a maximum e-value of 1e-5.

### Relative order of the acquisition events

We considered as main HGT donors for *P. micropora*, those taxonomic groups identified as donors in more than 10 HGT-C events. For each set of HGT-C proteins from the same main donor, we calculated the raw stem length, which is the length of the branch connecting the *P. micropora* sequence or family to the first sister clade. We obtained the normalised stem length, a relative measure of evolutionary time (Susko et al. 2021), by dividing the raw stem length by the median root-to-tip distance, following a procedure similar to Pittis and Gabaldón (2016).

## Data and code availability

Data and the archived code to produce the results of this study are available in a Zenodo repository upon publication.

## Acknowledgements

TG group acknowledges support from the Spanish Ministry of Science and Innovation (grant numbers PID2021-126067NB-I00 and PLEC2023-010225) cofounded by ERDF “A way of making Europe”, as well as support from the Gordon and Betty Moore Foundation (grant number GBMF9742); the Catalan Research Agency (AGAUR) (grant number 2022 INNOV 00065, 2024 PROD 00175 and 2024 PROD 00043); “La Caixa” foundation (grant number LCF/PR/HR21/00737, LCF/PR/HR21/52410006, LCF/PR/GN18/50310010 and CI23-20260); Fundació La Marató de TV3 (202328-31); AECC (PRYGN234923GABA and 290059); Instituto de Salud Carlos III (CIBERINFEC CB21/13/00061-ISCIII-SGEFI/ERDF and DTS25/00141); European Commission, Horizon Europe-HORIZON-MSCA-2023-DN-01-01 (grant number 101168618) and European Union’s Horizon 2020 research and innovation programme under the Marie Skłodowska-Curie grant agreement Nº 101226544 (grant number 101227078). We also acknowledge the CGenomics members for their valuable comments and help. We especially want to thank Giacomo Mutti for his help in the implementation of the annotation pipeline.

## Supplementary material content

Supplementary Methods, Results and Discussion, Figures 1-9 and References

## Supplementary tables content

Supplementary Tables 1-5.

