## Supplementary text and figures for "Host retargeting predominated over gene transfer during early chromatophore integration in *Paulinella micropora*"

**Supplementary materials index**

### Supplementary methods

#### ***Functional annotation pipeline***

To functionally annotate *P. micropora* proteins, we used EggNOGmapper v2.1.12 (Cantalapiedra et al. 2021) and InterProScan v5.57-90.0 (Jones et al. 2014), and performed profile-based searches using hmmsearch v3.3.2 (Eddy 2011) against the following databases: Database of Clusters of Orthologous Genes (COG) v2020 (Tatusov et al. 1997; Galperin et al. 2015), PFAM v35.0 (Mistry et al. 2021), KEGG Orthology (KO) v2023-02-01 (Kanehisa et al. 2016) and TIGR (Haft et al. 2013), which we downloaded on 20 April 2023. We filtered and merged these results into a single annotation file using a custom script (Supplementary Figure 6, see Data and Code availability). This script parses the hmmsearch results for each database to retrieve the first hit, and merges this information with the description of the HMM profile IDs, creating a table that relates the first hit to the description in the corresponding database (COG, KO, TIGR and PFAM). Subsequently, the script parses the tabular outputs of InterProScan, getting the description and the method. Finally, it reads the EggNOGmapper output, retrieving the description, the preferred name, the COG category and the gene ontology (GO) terms. Finally, for each protein, the script iterates through the annotations assigned by the different methods, and assigns one single annotation from a preferred method following this order: EggNOGmapper, InterProScan (excluding those hits from MobiDBLite and Coil databases), COG database, KO database, PFAM and TIGR.

**Supplementary results and discussion**

***Benchmarking of the HGT detection pipeline using the chromatophore genome***

To assess the performance of the HGT detection pipeline, we ran it over the proteins encoded in the chromatophore's genome, which has a known bacterial origin. For this, we annotated the chromatophore proteins as belonging to eukaryote and assessed the potential phylogenetic origins at the Phylum and Order levels. The pipeline correctly inferred non-eukaryotic origins for all chromatophore proteins except two (2 out of 859, 0.23%). For five proteins no origin at the Phylum level could be established due to the most abundant Phylum in the sister group representing less than 80% of the sequences (Supplementary Figures 2a, 3a). For the remaining proteins, a Phylum-level donor was found under the relaxed criterion in 836 trees at (97.32% of the genome), and 793 trees at the Order level (92.32% of the genome). When the strict criteria were applied, we found 794 trees with a specific Phylum-level donor found (92.43% of the genome), and 594 at the Order-level (69.15% of the genome).

As compared to the results obtained for the nuclear-encoded proteins (see main text), the taxonomic distribution is rather narrow, with only three phyla and five orders under the strict criteria. As expected from the known cyanobacterial origin for of the chromatophore, we detect few potential cases of transfer from other bacteria besides, four acquisitions from Proteobacteria and one from Actinobacteria at the Phylum level (Supplementary Figure 2b). At the Order level, the main detected phylogenetic origin is the PCC-6307 clade, which belongs to the Prochlorococcus-Synechoccocus clade, which is the known sister-clade of the chromatophore in the species tree. Two proteobacterial acquisitions, one from Rickettsiales and one from Burkholderiales, are detected at the Order level under the strict criteria. These results validate our HGT detection pipeline and shed light on the origin of the genes encoded by the chromatophore genome, indicating that horizontal transfers to the crhomatophore genome was minimal, with only five robust contributions from two different phyla (Supplementary Figure 2b, HGT-C).

***Taxonomic bootstrap analyses***

To assess the robustness of the inference of the taxonomic assignment of the donor, we designed a support measure at the Phylum level, rather than at the sequence level. That is, instead of assessing the robustness with regard to the specific sequences present in the sister-clade, we assessed the robustness of the taxonomic assignment derived from the Phylum affiliation of the sequences in the sister clade, as inferred by our pipeline. That is, we considered a high robustness in the assignments if the taxonomic inference remained consistent, regardless of variations on the specific set of sequences within the sister-clade. The logic behind this approach is that, given the long time since the transfers, we expect the phylogenetic signal to be blurred lacking specificity for the specific sister sequences, but we expect a higher reliance on the taxonomic affiliation at higher taxonomic levels.

For each family, we calculated the proportion of the most abundant taxa (the donor) in the sister groups to the *P. micropora* family both in the maximum likelihood (ML) tree and in the 1000 ultrafast bootstrap (UFBoot) trees. We defined as "taxonomic bootstrap support" the fraction of bootstrap trees that was consistent with the taxonomic affiliation (at the Phylum level) with the one derived from the ML tree topology. We noted that the number of families for which the most frequent donor in the UFBoot trees coincides with the ML donor outnumbers those in which they do not coincide (Supplementary Figure 7a). To test how the UFBoot bootstrap and the newly derived taxonomic bootstrap compare, we plotted the correlation between the two measurements (Supplementary Figure 7b). We observe that both measures correlate for most of the trees. The trees in which the ML and taxonomic bootstrap donors do not coincide are almost always located in the lower half of the UFBoot values. The cause of the trees in which we observe full UFBoot support values, but lower taxonomic bootstrap values, are due to rooting differences between the bootstrap tree and the ML tree. However, for another fraction of trees we observed high taxonomic bootstraps despite low UFBoot bootstrap values. These were caused by instability in the sequences contained within the sister clade, which nevertheless were always supporting the same taxonomic affiliation. Hence, we concluded that the taxonomic bootstrap method was more appropriate to assess the level of certainty of the taxonomic assessment.

***Evolution of the P. micropora chromatophore proteome***

We inferred the chromatophore's proteome by combining proteins encoded in the chromatophore genome (859, 75%) with the host-encoded and chromatophore-targeted (imported) proteins (287, 25%). This resulted in a proteome estimate of 1146 proteins, which is a significantly reduced proteome, as compared to the typical size of free-living relatives, with around 2500 proteins (Lhee et al. 2019). Using the results of the evolutionary analyses of the gene families, we assessed the origins of the chromatophore proteome. As expected, most proteins (848, 74%) have cyanobacterial origins (Figure 5, left panel), followed by vertically inherited genes encoded in the *P. micropora* nuclear genome and retargeted to the organelle (231, 20%). A set of 47 (4.1%) chromatophore proteins were horizontally acquired from other bacteria, of which 11 (1%) are encoded in the chromatophore genome, and 36 are nuclear-encoded (3.1%) genomes. These HGT genes have various origins, among which Gammaproteobacteria and Actinobacteriota are the most commonly inferred donors (Figure 4 and Figure 3b). Finally, we could not properly determine the origin of two chromatophore proteins. Overall the proteome's origin distribution reflects the major origin of the organelle's and host's genomes, with chromatophore-encoded proteins being mostly of cyonabacterial origins (Figure 5, centre), and nuclear-encoded imported proteins mainly evolving vertically from the host (Figure 5, right). Functionally, the only functional category with overrepresented non-cyanobacterial HGTs is replication, recombination and repair (12). hese genes are: DNA ligase, several ATP-dependent RNA and DNA helicases, DNA repair protein RecN, ATP-dependent helicase HrpB and the exodeoxyribonuclease V alpha subunit.

***Expanded P. micropora chromatophore's imported proteome***

*Paulinella* species have two types of proteins imported into the chromatophore: the short imported peptides (sCTPs) and the long chromatophore targeted peptides (lCTPs). The lCTPs contain an N-terminal import signal, the chromatophore transit peptide (crTP), which has approximately a length of 200 amino acids (Singer et al. 2017). The crTP sequence was used in Lhee et al. (2021) to identify the lCTPs in *P. micropora*, which is the set of chromatophore proteins we used in our inferences. To expand this set and include sCTPs in *P. micropora*, which cannot be predicted bioinformatically, we retrieved the sequences of the 433 *P. chromatophora* experimentally-determined import candidates in Singer et al. (2017), and performed a blast-based homology search of these proteins in the inferred proteome of *P. micropora*.

We found homologues for all the import candidates, and detected an expansion of the "scaffold63687-m.206362" protein, which has 11 identical copies in *P. micropora*, and is annotated as high-light-inducible (HLI) protein. These results agree with the already reported frequent expansions of HLI, likely mediated through retrotransposition (Calatrava et al. 2022), and important for the adaptation to high-light stress. For the rest of the import candidates, we only considered the first Blast hit.

The distribution of coverage and identity of the first hits (Supplementary Figure 9 and Supplementary Table 2) reveals a mean identity around 50% with a minimum threshold of about 30%. Query coverage distribution shows a mean coverage of 76% for the first hits. Despite these average values, the distribution reveals high uncertainty for the hits. The coverage distribution is left-skewed, and 42% of the hits have a coverage lower than 75%. In contrast, the identity distribution is right skewed, with only 11% of the data showing values higher than 66% identity. The identity mode is around 30% (a value generally assumed as a minimum threshold to detect homologues). Additionally, our previous search of *P. micropora* homologues in *P. chromatophora* discarded most of these hits as they did not fit our thresholds, despite these being already quite relaxed (see Methods). Overall, although protein similarity is related to function and function to localisation (Nair and Rost 2002), these results evidence that assigning sCTPs in P. micropora using sequence similarity to experimentally-determined sCTPs in *P. chromatophora* is difficult, as we can not trace back one-to-one relationships between these proteins. Therefore, we decided not to use this expanded set in the primary analyses.

Nevertheless, to test the impact that including these sCTPs homologues would have in our conclusions, we annotated as proteins localised in the chromatophore the first hits to the import candidates in Singer et al. (2017), resulting in a set of 443 CTPs. 256 of these correspond to the lCTPs previously assigned to the chromatophore using Lhee et al. (2021), and the remaining 187 are putative sCTPs derived only from the homology search. Using these expanded dataset of proteins imported in the chromatophore, the main conclusions drawn by the lCTPs are maintained. Vertically inherited proteins remain more frequent among the imported proteins (14% in the expanded dataset and 13% among the lCTPs, Supplementary Figure 4b). However,  the use of the expanded dataset increased the number of horizontally transferred genes from cyanobacterial donors (Supplementary Figure 4c). This agrees with previous results in *P. chromatophora* (Nowack et al. 2016) and *P. micopora* (Lhee et al. 2021), where cyanobacterial well-known putative import candidates (i.e., EGTs) are also discussed.

The functional profile of the imported genes separated by their origin shows that most functional categories are dominated by vertically inherited genes (Supplementary Figure 4d). However, the "Replication, recombination and repair (L)" category presents a higher number of proteins from horizontal origin in the lCTPs set (Figure 4d). Conversely, in the expanded dataset their differences are smaller, strengthening the conclusions that overall, the vertically inherited proteins targeted to the chromatophore dominate most of the chromatophore's functions.

***Metabolic integration of targeted genes in the chromatophore***
To explore how nuclear-encoded ICTPs targeted to the chromatophore are integrated in the metabolic network of *P. micropora*'s chromatophore, we collected KEGG ortholog (KOs) annotations for the genes expressed in the chromatophore. These genes have three origins: 1) the chromatophore's genome, 2) nuclear-encoded from a vertical origin, and 3) nuclear-encoded with an HGT origin.

To compare the current chromatophore's proteome with the putative ancestral one, we reconstructed a proxy of the ancestral chromatophore's genome using a pangenome approach of the closest species to the chromatophore. For this, we selected 7 reference genomes from NCBI for the Synechococcus / Prochlorococcus clade that appears sister to the *Paulinella* genus chromatophore according to Lhee et al. (2019). Then, we clustered the 7 proteomes using mmseqs easy-cluster with default parameters (0% minimum identity, and 80% minimum coverage), and kept for the pangenome those present in both clades (Synechococcus, and Prochlorococcus) or was present in at least 3 species out of 7. This resulted in a Prochlorococcus / Synechococcus pangenome of 2,105 proteins. We annotated the reference proteome using KOFamScan v1.3.0 (Aramaki et al. 2020). When a significant KO was present, we assigned it to the protein, otherwise, we kept the first hit as the closest annotation. This constituted the "ancestral" set of KOs. In total, we annotated 1451 proteins, 620 of them as significant hits (in the zenodo data repository we share the list of used genomes, the pangenome, and their functional annotation).

We used KEGG mapper color (https://www.genome.jp/kegg/mapper/color.html) to highlight enzymes present in each set in all the available metabolic pathways (see the data repository). Although these metabolic maps are biased towards model organisms, and they may not represent the actual system in the chromatophore, they are valuable to observe regions of these pathways present in the chromatophore.

Similarly to what we observe in the functional enrichment analysis (Figure 3), the metabolism and transport of amino-acids, nucleic acids and lipids are the maps that show more complete pathways. We find complete isoleucine, valine, and leucine biosynthesis pathways (Supplementary Figure 5a). In these three pathways, which share some enzymes, we find that all the genes present in the putative ancestral proteome are also found in the chromatophore's genome, except for the 3-isopropylmalate dehydrogenase [EC:1.1.1.85] (K00052). This gene is present in the proxy of the ancestral proteome and in a nuclear-encoded gene that is targeted to the chromatophore. This suggests that the ancestral gene was replaced with a targeted gene. Similarly, the synthesis of proline from glutamate shows the same pattern. The first steps of the pathway (catalysed by the glutamate 5-kinase –K00931–, and the glutamate-5-semialdehyde dehydrogenase –K00147) are present in the genome of the chromatophore and the putative ancestral one. However, the final step (catalysed by pyrroline-5-carboxylate reductase –K00286), is present in the putative ancestral pangenome and the nuclear genome and targeted to the chromatophore, suggesting another replacement (Supplementary Figure 5b). This trend becomes even more general in other pathways like in purine and pyrimidine metabolism (see the data repository for a detailed view of the pathways).

Regarding HGT-derived nuclear-encoded genes that are targeted to the chromatophore, we see a scattered pattern across the pathways. This suggests that HGT genes targeted to the chromatophore may have contributed to broadening the metabolic network of the chromatophore with new metabolic capacities that were likely absent in the ancestor , rather than replacing functions lost during the endosymbiotic ratchet. To test this hypothesis, we additionally ran anvi-estimate-metabolism, from the anvi'o suite v9 (Eren et al. 2021). We filtered out incomplete pathways according to the default stepwise completeness (> 75%), and inferred the origins of each step (Supplementary Table 3 and the data repository). In summary, we find that among the complete pathways, we observe that targeted genes that evolved vertically in the host substituted the ancestral ones in 8 different module steps. Conversely, the substitution of ancestral genes for HGT targeted genes only happens once. This trend also applies to the genes that were not present in the chromatophore's ancestor, which would suggest pathway steps introduced by targeting (targeting-derived "innovations"). Host's vertical genes incorporated 5 new steps through targeting into the chromatophore, whereas HGT shows only one.

Taking all these observations together, the metabolic reconstruction of the chromatophore shows a higher impact of substitutions of ancestral proteins with targeted ones that evolved vertically in the endosymbiotic host. Similar to what we observe in the proportion of the origin of targeted genes when adding putative sCTPs, we would expect similar trends of completeness and metabolic complementation due to substitution of steps with vertically inherited genes in the host. However, although we acknowledge the incompleteness of our reconstruction due to the absence of reliable sCTPs, we cannot risk making robust and reliable metabolic nor evolutionary inferences on a homology-based inferred set of sCTPs.

### Supplementary figures

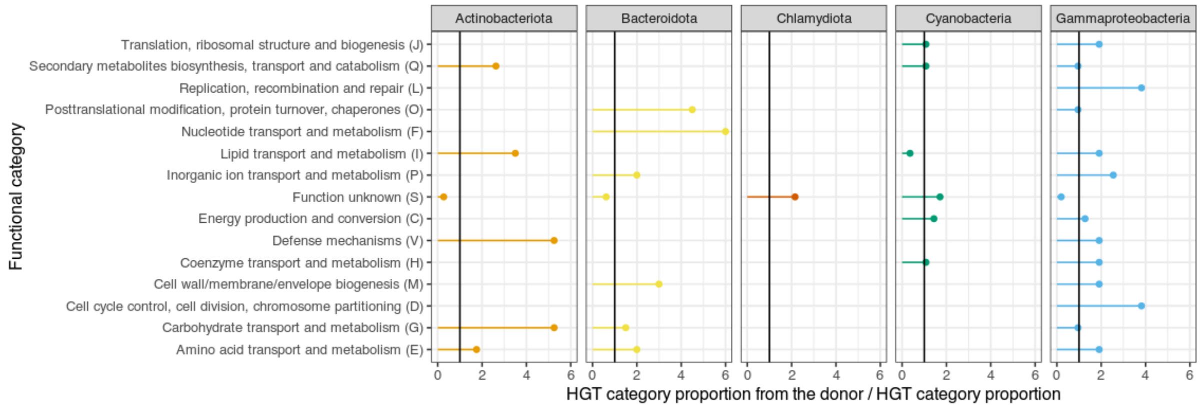

**Supplementary Figure 1. Enriched functions in the main donors**. Each point shows the value obtained by dividing the proportion of genes annotated with a category among HGT genes from each donor by the overall proportion of the same category in the whole set of HGT genes. Values above one (black solid vertical line) show a quantitative enrichment of the function in the HGTs from the donor. We did not use hypothesis contrast due to small sample sizes, Hence, these values should be interpreted carefully.

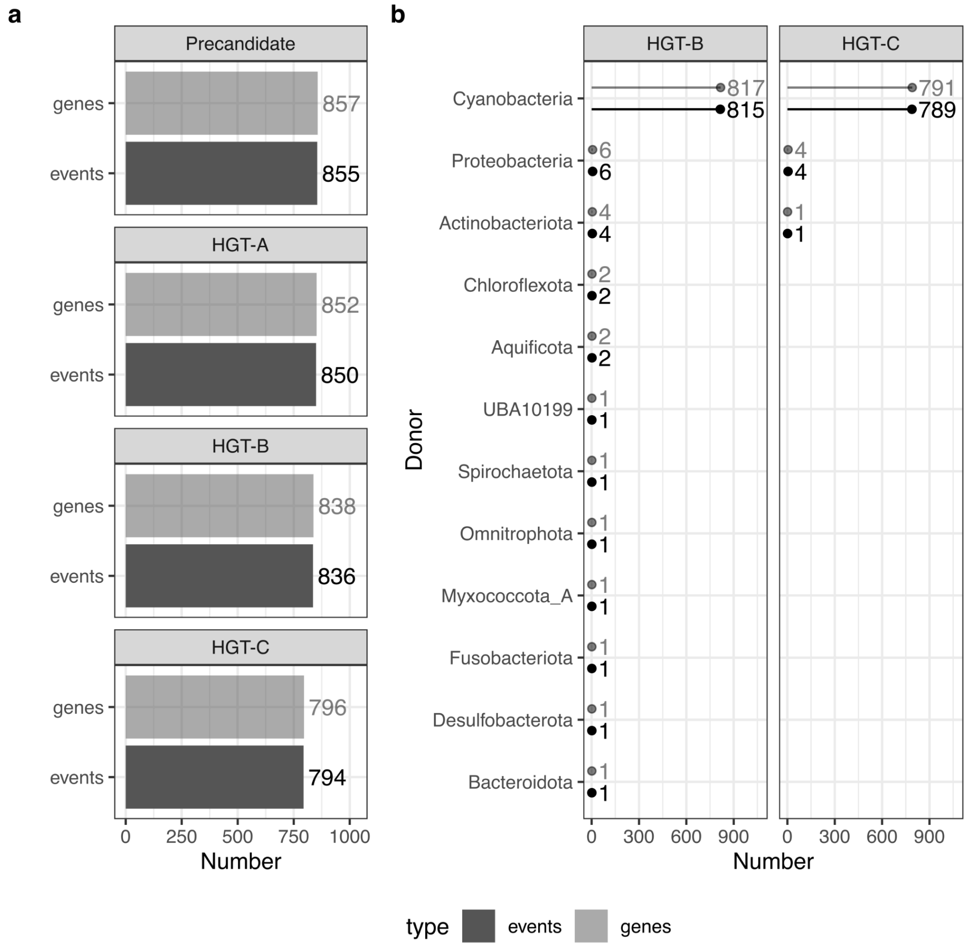

**Supplementary Figure 2. Horizontal gene transfer analysis with the chromatophore genome at the Phylum level**. a) Count of genes and events in each step of the pipeline as in the main Figure 1. b) Taxonomic distribution at the Phylum level of the organisms involved in the detected HGT. Note that in this case the cyanobacterial genes would not be acquisitions but vertical evolution, as the working genome evolved from cyanobacteria.

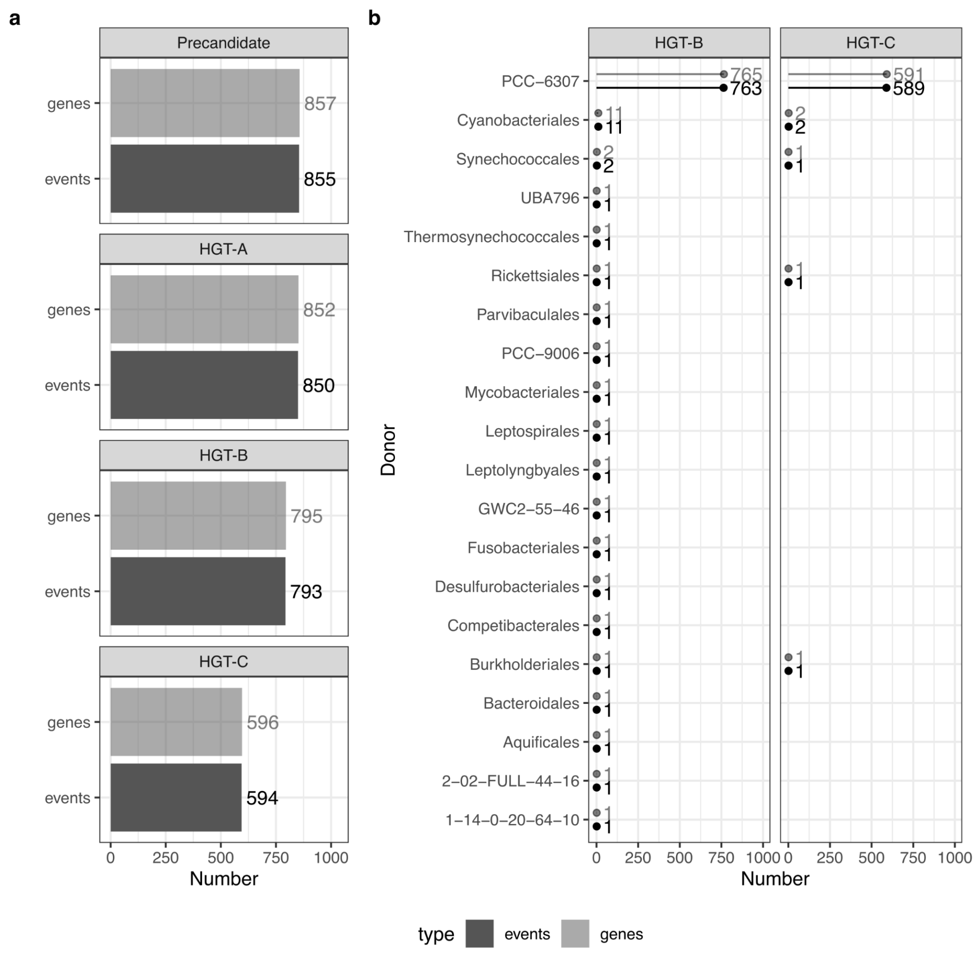

**Supplementary Figure 3. Horizontal gene transfer analysis with the chromatophore genome at the Order level**. a) Count of genes and events in each step of the pipeline as in the main Figure 1. b) Taxonomic distribution at the Order level of the organisms involved in the detected HGT. Note that in this case the cyanobacterial genes would not be acquisitions but vertical evolution, as the working genome evolved from cyanobacteria.

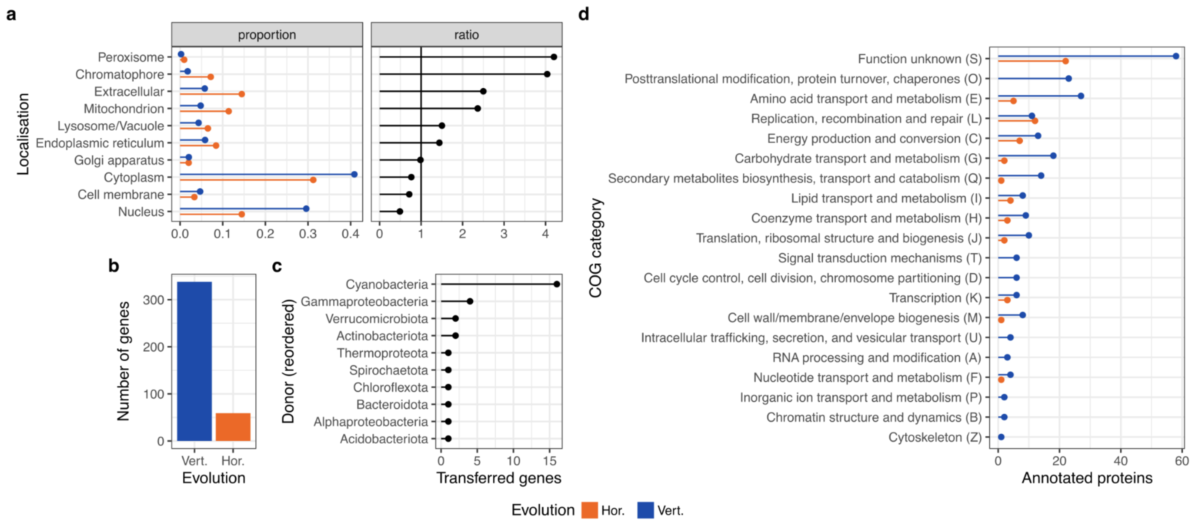

**Supplementary Figure 4. Subcellular localisation of the *P. micropora* proteins based on chromatophore target peptides (crTP) and homologues to the imported peptides in Singer et al. (2017)**. a) Proportion of proteins from each origin assigned by DeppLoc to each subcellular compartment, and the corresponding proportion ratio. Ratio values greater than 1 indicate that the predicted localisation is more frequent among HGT-derived proteins than among vertically-inherited proteins. b) Number of nuclear-encoded proteins predicted to be targeted to the chromatophore based on the presence of a chromatophore transit peptide (crTP) as inferred by Lhee et al. (2021), grouped by inferred origin (vertical or horizontal). c) Putative donor of the HGT-derived proteins predicted to be targeted to the chromatophore. d) Functional categories of chromatophore-targeted proteins according to inferred origin.

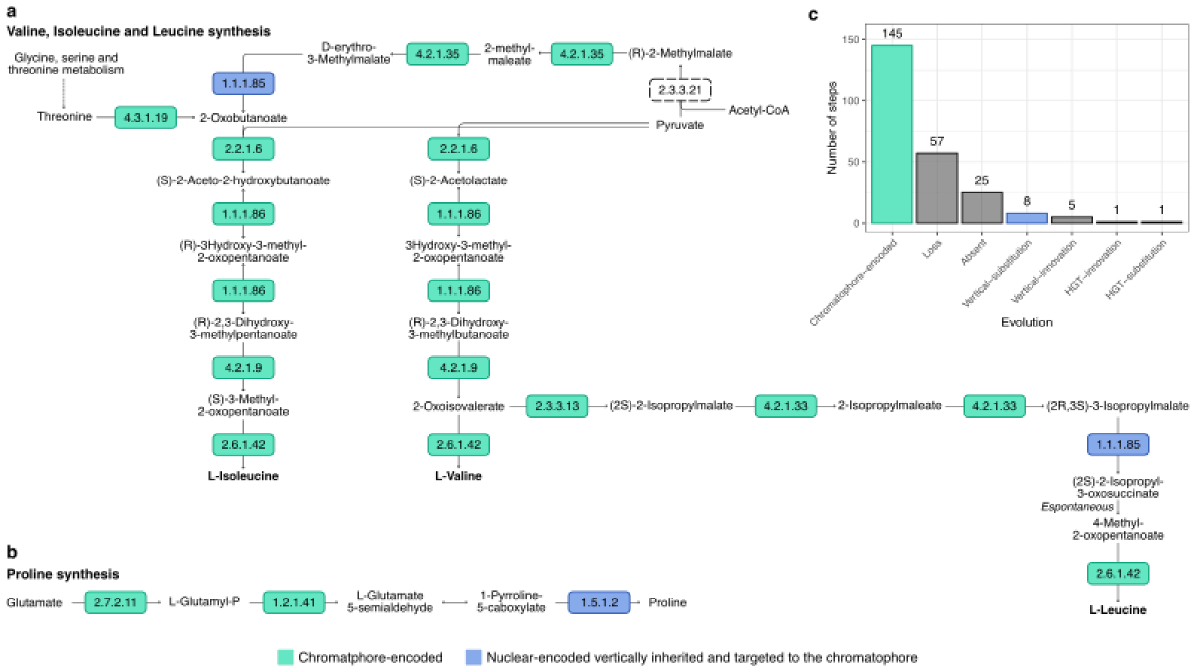

**Supplementary Figure 5. Integration of host's vertically inherited genes in the chromatophore metabolism**. a) Pathways for the synthesis of valine, isoleucine and leucine. b) Pathway of the Proline synthesis. c) Number of steps of all the complete pathways according to the origins of their enzyme. The colours of the enzyme boxes and the bars in c) corresponds to the origin accoridng to the palette in the bottom.

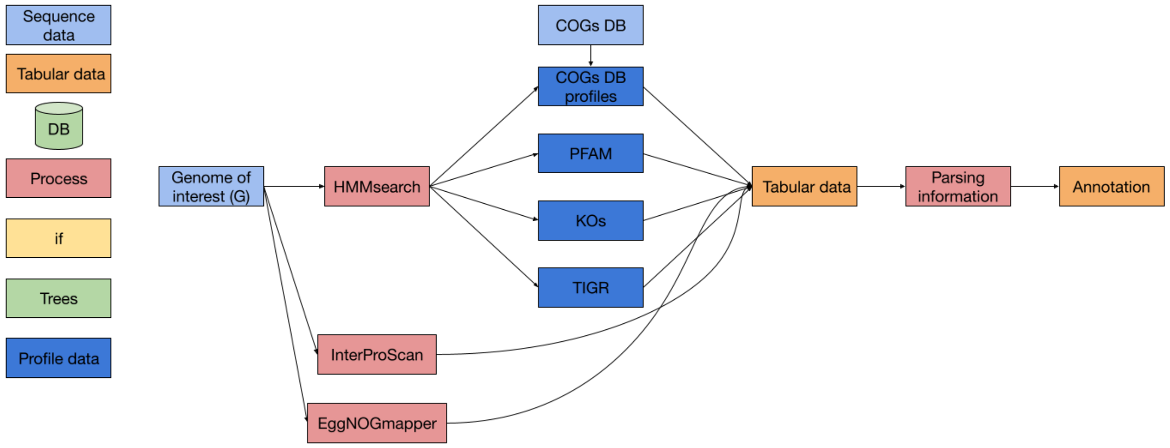

**Supplementary Figure 6. Annotation pipeline scheme**. The blocks on the left show the colour and shape of the different kinds of steps.

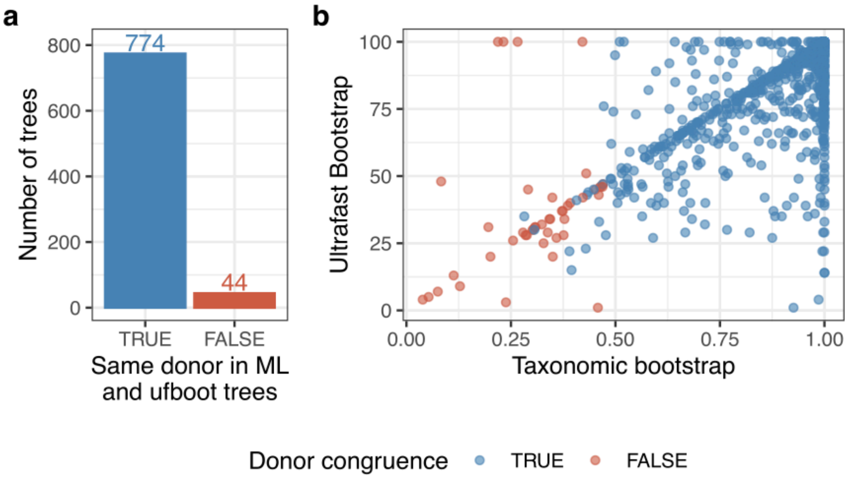

**Supplementary Figure 7. Taxonomic bootstrap analysis**. a) Number of trees in which the donor inferred in the ML tree and the most abundant clade inferred as donor in the ultrafast bootstrap trees are congruent (true) or not (false). b) Correlation between the ultrafast bootstrap values for the branch separating the *P. micropora* family and its first sister and the support of the ML donor in the bootstrap trees.

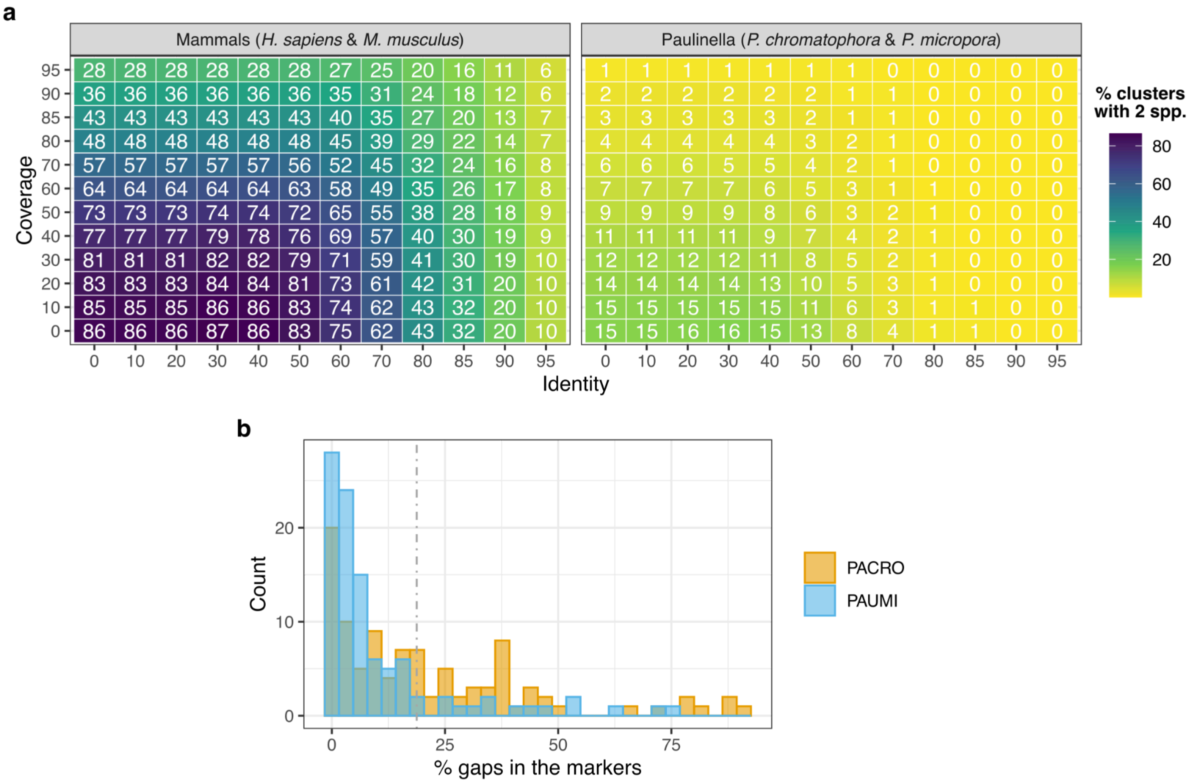

**Supplementary Figure 8. Analysis of the completeness of *P. chromatophora***. a) Heatmap of the percentage of sequence clusters with members of both species indicated in the panel title according to different identity and coverage thresholds. b) Histogram of the percentage of gaps in the alignment of BUSCO genes found in *P. chromatophora* (PACRO) and *P. micropora* (PAUMI), shown in colours according to the legend. The vertical line shows the 20% of the gaps.

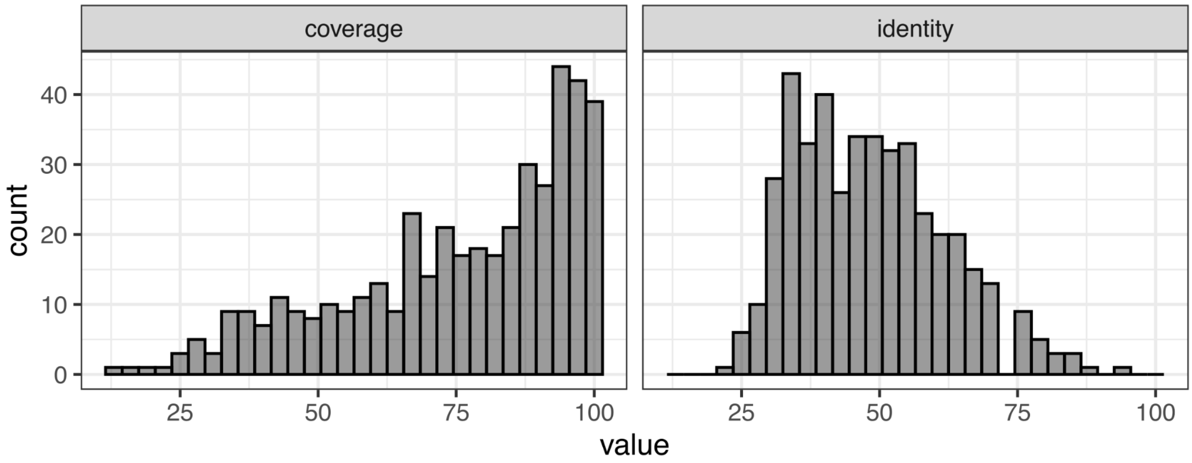

**Supplementary Figure 9. Similarity parameters for the *P. chromatophora* import candidate homologues in *P. micropora***. Distribution of the coverage (left) and identity (right) for the first hits of all the import candidates found in *P. micropora*.

### Supplementary tables

**Supplementary Table 1. Summary of the annotation, localisation and evolutionary origin of the nuclear-encoded and chromatophore-encoded *Paulinella micropora* proteins**. The localisation colum are DeepLoc predictions, a deep learning algorithm to predict subcellular localisation. The accuracy of the method is poor for not well-described organisms, therefore we warn the reader that this results should be interpreted cautiously. When the Localisations value takes "Chromatophore" the protein corresponds to the crTP targeted proteins inferred in Lhee et al. (2019).

**Supplementary Table 2. *Paulinella chromatophora* import candidate homologues in *P. micropora***. The proteins that lack the columns from "Family reference" to "COG cagtegory" is because they do not have functional annotation.

**Supplementary Table 3. Origins of the complete pathways in the chromatophore proteome**. Completeness is assessed with Anvi'o, which takes the stepwise-completeness > 75% as a threshold.

**Supplementary Table 4. OMArk completeness statistics**.

**Supplementary Table 5. OMArk contamination statistics**.

### Data and code availability

Data and the archived code to produce the results of this study are available in a Zenodo repository upon publication.
